# The soluble N-termini of mechanosensitive ion channels MSL8, MSL9, and MSL10 are environmentally sensitive intrinsically disordered regions with distinct biophysical characteristics

**DOI:** 10.1101/2022.10.14.512244

**Authors:** Aidan J. Flynn, Kari Miller, Jennette M. Codjoe, Matthew R. King, Ammon E. Posey, Elizabeth S. Haswell

**Affiliations:** Department of Biology, Washington University in St. Louis, St. Louis, MO, USA; NSF Center for Engineering Mechanobiology, Washington University in St. Louis, St. Louis, MO, USA; Department of Biochemistry and Biophysics, Washington University in St. Louis, St. Louis, MO, USA; Department of Biomedical Engineering, Washington University in St. Louis, St. Louis, MO, USA

**Keywords:** Arabidopsis, mechanobiology, transmembrane protein, ion channel, intrinsically disordered protein, circular dichroism, phase separation

## Abstract

Intrinsically disordered protein regions (IDRs) are highly dynamic sequences that rapidly sample a collection of conformations. In the past several decades, IDRs have emerged as a core component of many proteomes, comprising ∼30% of all eukaryotic protein sequences. IDRs are ubiquitous throughout different biological pathways, with a notable enrichment in responses to environmental stimuli such as abiotic stress. However, the diversity of IDR-based systems that biology has evolved to respond to different stimuli is expansive, warranting the exploration of IDRs present in unique molecular contexts. Here, we identify and characterize intrinsic disorder in the soluble, cytoplasmic N-terminal domains of three members of the MscS-Like (MSL) family of mechanosensitive ion channels, MSL8, MSL9 and MSL10. In plants, MSL channels are proposed to mediate the reactions to cell swelling, pathogenic invasion, and touch. A series of bioinformatic tools unanimously predicted that the cytosolic N-termini of MSLs are intrinsically disordered. We confirmed this prediction for the N-terminus of MSL10 (MSL10^N^) via circular dichroism spectroscopy. MSL10^N^ adopted a predominately helical structure when exposed to the helix-inducing compound trifluoroethanol (TFE) and underwent structural changes and alterations to homotypic interaction favorability in the presence of molecular crowding agents. Lastly, *in vitro* imaging of condensates indicated that MSL8^N^, MSL9^N^ and MSL10^N^ have sharply differing propensities for condensate formation both inherently and in response to salt, temperature, and molecular crowding. Altogether, these data establish the N-termini of MSL channels as intrinsically disordered regions with distinct biophysical properties and the potential to respond disparately to changes in their physiochemical environment.

## INTRODUCTION

In recent years, the function of intrinsically disordered regions (IDRs) in plant biology has become an attractive topic due to their putative roles in myriad functions, including transcription, scaffolding, and stress responses (Covarrubias et al., 2017; Emenecker et al., 2020). IDRs are protein regions that lack rigid three-dimensional structure, instead dynamically sampling a series of conformations over time known as a conformational ensemble (Dunker et al., 2013). These conformational ensembles sometimes dictate the biological function of the disordered region and are often influenced by the surrounding cellular context (Das and Pappu, 2013; Mao et al., 2012; Cohan et al., 2019). Due to this environmental sensitivity, IDRs are proposed to provide a molecular mechanism for sensing and responding to intracellular stress (Cuevas-Velazquez and Dinneny, 2018; Emenecker et al., 2020). Further, several natural IDRs have been repurposed for use as environmental sensors in experimental settings (Cuevas-Velazquez et al., 2021).

IDRs localize to a variety of different cellular contexts and compartments. Yet, only a fraction of IDRs present at membranes have received detailed experimental characterization (Kjaergaard and Kragelund, 2017; Verkest et al., 2022; Goretzki et al., 2021), despite a significant enrichment of IDRs in cytosolic extensions of membrane proteins compared to the full proteome (Bürgi et al., 2016). Because of the established importance of IDRs in environmental sensing and signaling and their proximity to the site of physical stress recognition (the membrane), membrane-tethered IDRs are an attractive model by which cells may rapidly recognize and respond to stress response.

Mechanosensitive (MS) ion channels are transmembrane channels that display a characteristic opening following mechanical force, thereby allowing ions to travel across the membrane. In plants, these channels are implicated in the response to a wide variety of stimuli, including germination, cell wounding, and osmotic stress (Basu and Haswell, 2017). In the model flowering plant *Arabidopsis thaliana*, there are ten members of the MscS-like (MSL) protein family that share homology with the *Escherichia coli* MS channel MscS (<u>M</u>echano<u>s</u>ensitive <u>c</u>hannel of <u>S</u>mall conductance) (Pivetti et al., 2003; Haswell, 2007). The ten MSLs in Arabidopsis have differing expression profiles and localize to various organellar membranes, including the mitochondria, plastid, and plasma membranes (Haswell, 2007; Lee et al., 2016; Haswell and Meyerowitz, 2006; Haswell et al., 2008).

Generally speaking, MSL channels are thought to act as “osmotic safety valves” that prevent organelle or cell bursting by opening in response to increased membrane tension (Basu and Haswell, 2017). However, additional functions, and modes of regulation, have been proposed (Wang et al., 2021; Veley et al., 2014). For example, MSL10 promotes programmed cell death when overexpressed or in response to cell swelling in seedlings (Veley et al., 2014; Basu et al., 2020; Basu and Haswell, 2020), perhaps as a response to cell wall damage or osmotic changes during pathogenic invasion (Basu et al., 2022). This activity is inactivated by phosphomimetic mutations at 7 known phosphorylation sites in the soluble 164-residue N-terminus of MSL10 (Veley et al., 2014; Basu et al., 2020; Basu and Haswell, 2020). The cell death promoting activity of MSL10 is both physically and genetically separable from its function as a mechanosensitive ion channel (Maksaev et al., 2018).

MSL10 is localized to the plasma membrane and expressed ubiquitously throughout the plant (Haswell et al., 2008). The channel has a slight preference for anions but is otherwise non-selective (Haswell et al., 2008; Maksaev and Haswell, 2012). MSL9, a close homolog of MSL10, is expressed in root tips and has no known genetic function. MSL9 and MSL10 likely form a mixture of homomeric and heteromeric complexes *in vivo* (Haswell et al., 2008). A third member of the MSL channel family, MSL8, is expressed in pollen where it is required for normal survival of pollen grain rehydration and pollen tube growth(Hamilton et al., 2015; Wang et al., 2021). Here, we provide an analysis of intrinsic disorder in the soluble N-termini of MSL8, MSL9, and MSL10, using both *in silico* and *in vitro* approaches. We find that these regions are predicted to be intrinsically disordered and demonstrate that this disorder is likely ubiquitous throughout putative orthologs of MSL9 and MSL10 in angiosperms. Our work collectively establishes the presence and implicates the relevance of IDRs in the functions of MSLs, while additionally reporting a new collection of evolutionarily related IDRs with notably different environmental responses.

## RESULTS

### The N-termini of plasma membrane-localized MSL channels have characteristics of intrinsically disordered regions

Of the ten MSL channels in Arabidopsis, MSL8-10 are predicted or shown to be localized to the plasma membrane and have the same predicted topology (Haswell, 2007). They are comprised of a soluble N-terminal domain of approximately 150 aa, six transmembrane helices, a soluble domain between TM helices 4 and 5, and a soluble C-terminal domain that includes the MscS homology domain which forms the pore of the channel (**Figure 1A**). There is low sequence conservation between Arabidopsis MSLs outside of the pore-lining and transmembrane domains, and this sequence divergence is especially visible in the soluble N-terminal domains of these proteins **(Figure S1A)**. As evolutionarily related IDRs often exhibit high sequence variability (Wallmann and Kesten, 2020; Zarin et al., 2017), this lack of conservation drove us to hypothesize that the MSL N-termini are intrinsically disordered.

**Figure 1.**
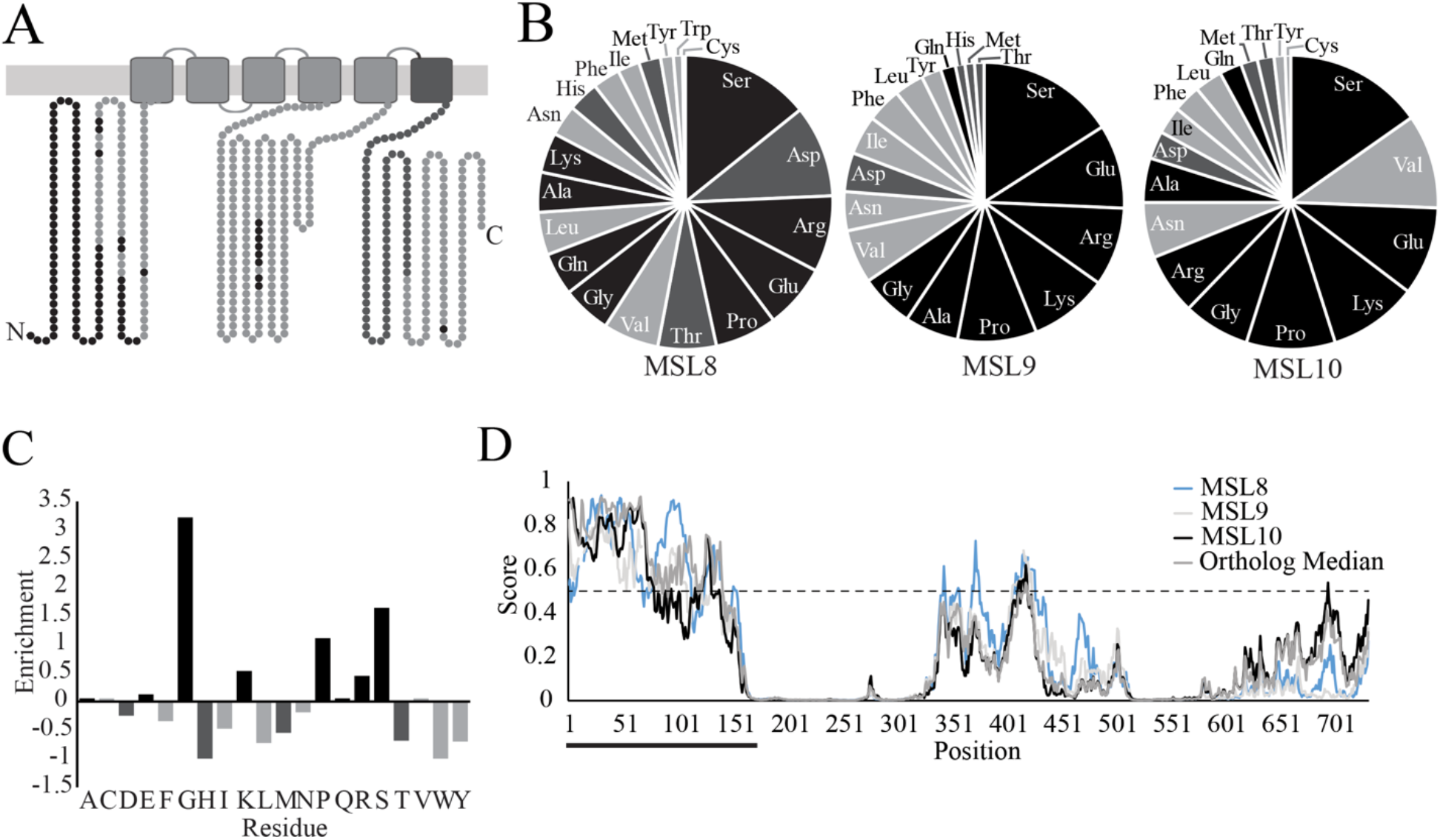
The N-termini of MSL family proteins are predicted to be disordered. (A) Topology and predicted disorder for MSL10. MSL8 and MSL9 present similar topologies. Residues predicted by IUPRED2A to be disordered arc highlighted in black. The conserved MscS domain of MSL 10 is dark grey. (B) Amino acid composition of the N-termini of MSL8, MSL9, and MSL 10. Calculations were performed using the PredictProlein webserver. Disorder-promoting residues are indicated with black, order-promoting residues with light grey, and disorder-order neutral residues with dark grey. (C) Compositional profile of the MSL10 N-tcrminus compared to the MSL10 C-tcrminus (aa 562-734). Comparisons were performed using the Composition Profiler webtool. Enrichmcnt of a particular amino acid is calculated as (Composition N-tcrminus - Composition C-tcrminus) / Composition C-terminus, where composition is the fractional amount of an amino acid within the N- or C-terminus. (D) IUPRED2A disorder profiled for full-length MSL8, MSL9, MSL10, and the median value of selected orthologs at a given aligned position of MSL 10. Residues with a disorder score higher than 0.5 arc predicted to be part of a disordered region, indicated by the dashed black line. Black bar indicates the N-tcrminal regions of each prediction.

Many IDRs are defined by characteristic amino acid compositions such as a depletion of hydrophobic residues that drive protein folding events and an enrichment in hydrophilic, charged, and polar residues (Uversky, 2013, 2019; Lee et al., 2014; Yachdav et al., 2014). We therefore assessed the amino acid frequencies present in the soluble N-termini of MSL8, MSL9, and MSL10. As shown in **Figure 1B**, the N-termini of MSL8 (amino acids 1-296, hereafter referred to as MSL8^N^), MSL9 (amino acids 1-176, hereafter referred to as MSL9^N^) and MSL10 (amino acids 1-164, hereafter referred to as MSL10^N^) contained enrichments of disorder-promoting residues like serine, glutamate, arginine, proline, and glycine, and depletions of order-promoting residues like cysteine, tyrosine, isoleucine, tryptophan, and phenylalanine. To further determine if this amino acid makeup is specific to the N-termini of MSLs, we compared the composition of exemplar MSL10^N^ to that of the structured MSL10 C-terminus through the Composition Profiler webtool (Vacic et al., 2007) **(Figure 1C)**. MSL10^N^ is significantly enriched in residues associated with disorder and depleted in order-promoting residues relative to the MSL10 C-terminus. Collectively, these results demonstrate that the N-termini of MSL8, MSL9, and MSL10 have sequence characteristics commonly associated with IDRs.

### The N-termini of MSL8-10 and homologs are predicted to be disordered

We next assessed the consistency of MSL IDR prediction by using a panel of web-server-based predictors, including IUPred2A, PONDR-VLXT, SPOT-Disorder2, PrDOS, and MFDp2. All algorithms used predicted a similar disorder profile in exemplar MSL10^N^ **(Figure S2A-D)**. Thus, we elected to use one algorithm, IUPRED2A, for the prediction of disorder in MSL8, MSL9, and MSL10 **(Figure 1D)** (Mészáros et al., 2018). MSL8^N^, MSL9^N^, and MSL10^N^ were predicted to be disordered, with MSL8^N^ having the highest propensity for disorder. The cytoplasmic loop (amino acids 306-512 of aligned profiles) was similarly predicted to be at least partially disordered for all three MSLs examined. Disordered regions are often highly variable in terms of primary sequence; however, the structural character of disorder within an IDR is sometimes highly conserved (Wallmann and Kesten, 2020). Our above findings indicates that intrinsic disorder is likely such a conserved feature within Arabidopsis MSLs. We also aligned the amino acid sequence of 13 putative orthologs of MSL9 and MSL10 from monocots and dicots (as reported in (Basu et al., 2020) and computed the median of their disorder scores at each position aligned with MSL10 (Christensen et al., 2019) **(Figure S1B, Figure 1D)**. The median score predictions closely matched the profiles of *A. thaliana* MSL9 and MSL10. These data cumulatively suggest that intrinsic disorder is an evolutionarily conserved property of the N-termini of plasma-membrane localized MSL channels and their homologs in monocots and dicots.

### The N-terminus of MSL10 is disordered *in vitro* and aggregates at high temperatures

To test our disorder predictions experimentally, we focused on MSL10, as its N-terminal domain has known functions *in vivo* (Veley et al., 2014; Basu et al., 2020; Maksaev et al., 2018)We first performed far-UV circular dichroism (CD) spectroscopy on purified C-terminally His-tagged MSL10^N^ **(Figure 2A)**. Measured spectra presented a negative peak near 200 nm (202 nm) that is typical of disordered proteins (Na et al., 2018) **(Figure 2B)**. Additionally, a slight shoulder was visible at 222 nm, which is indicative of some residual helical structure within the N-terminus (Greenfield, 2006). When we subjected purified His-tagged MSL10^N^ to increasing temperature, the negative peak at 202 nm gradually reduced in magnitude but did not shift left or right **(Figure 2B)**. This behavior is associated with soluble aggregate formation via homotypic interactions (Urry and Ji, 1968). Observed aggregation due to increased temperature is likely limited, as no substantial changes were seen in the HT voltages associated with CD spectra captured at each temperature **(Figure S3)**. Thus, the N-terminus of MSL10 is largely disordered and may experience an increased tendency for homotypic interactions and soluble aggregation with increasing temperature.

**Figure 2.**
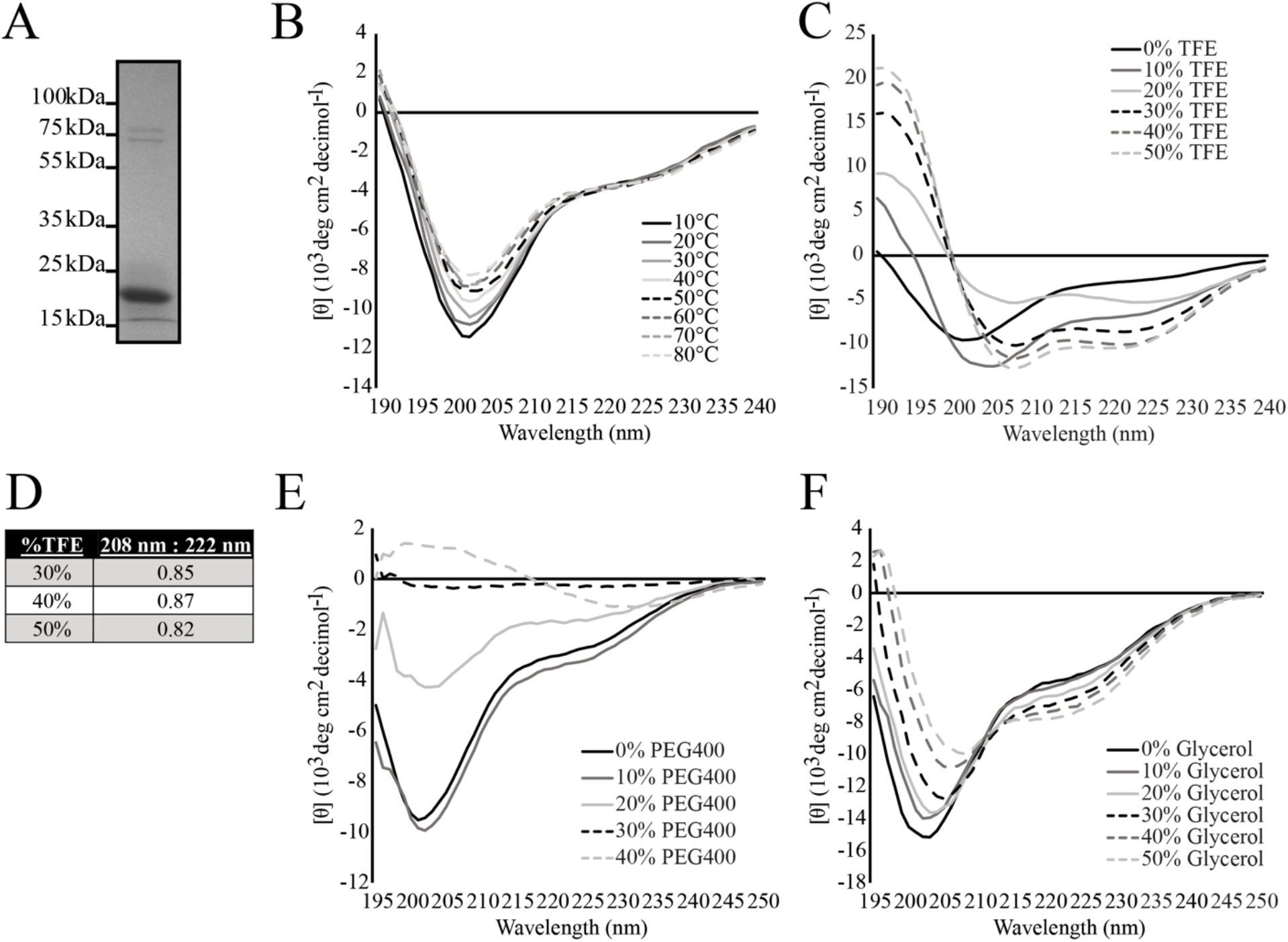
The MSL10 N-terminus is disordered and structurally responds to a variety of environments in *vitro*. (A) Representative coomassie-stained 10% SDS-PAGE gel of His-tagged MSL10^N^. His-tagged MSL10^N^ has an approximate molecular weight of 19 kDa. (B,C E, F) Circular dichroism spectra of His-tagged MSL10^N^when exposed to increasing temperatures (B), TFE (C), PEG 400 (D), and glycerol (E). Spectra were obtained at 20°C in 20 mM sodium phosphate buffer, pH 7.4 unless otherwise specified. (D) The ratio of the mean residue ellipticity value at 222 nm and 208 nm for 30%-50% TFE treatment.

### The N-terminus of MSL10 undergoes structural changes when exposed to helical-inducing compounds *in vitro*

The N-terminus of MSL10 has a relatively low fraction of charged residues, resulting in its classification as a Janus sequence on the Das-Pappu plot **(Figure S2E)**. IDRs that fall under this classification may take on coiled or globular ensembles depending on environmental contexts such as protein interaction (Das and Pappu, 2013; Das et al., 2015). To further examine the potential of MSL10^N^ to participate in protein interactions that cause disorder-to-order transitions, we introduced 2,2,2-Triflouroethanol (TFE) to His-tagged MSL10^N^ and measured structural changes via CD. TFE induces structure in disordered regions with a susceptibility to forming alpha-helices, such as for those that fold upon protein interaction (Luo and Baldwin, 1997; Hua et al., 1998; Chemes et al., 2012). As shown in **Figure 2C**, the addition of TFE resulted in the loss of the negative peak at 202 nm and the establishment of two negative peaks at approximately 208 nm and 222 nm. This profile is characteristic of helical proteins (Greenfield, 2006). For MSL10^N^ in 30%-50% TFE, the ratio of values at 222 nm to 208 nm was less than 0.9 **(Figure 2D)**. A value of less than 0.9 for this ratio signifies that the acquired helicity of MSL10^N^ occurs in an isolated helix form, as opposed to the coiled-coil structures associated with ratiometric values greater than 1.10 (Lau et al., 1984). These findings establish that the MSL10 N-terminus is capable of folding in the presence of TFE *in vitro*.

### MSL10’s N-terminus is sensitive to molecular crowding *in vitro*

Previous *in vitro* studies have demonstrated changes in plant IDR structure in response to molecular crowders and small molecules in physiologically relevant contexts (Dorone et al., 2021; Rivera-Najera et al., 2014; Mansouri et al., 2016; Chemes et al., 2012). We therefore conducted CD with His-tagged MSL10^N^ in the presence of polyethylene glycol 400 (PEG 400), a large polymer that induces simplified crowding conditions (Nakano et al., 2014), and glycerol, a small molecule that similarly simulates crowding (Rivera-Najera et al., 2014; Cuevas-Velazquez et al., 2016). In response to 10% and 20% PEG 400, the disordered minimum at 202 nm shifted rightwards **(Figure 2E)**. At 20% PEG 400, a significant flattening of the spectrum was also observed. Measurements in 30% PEG 400 resulted in near-zero absorbances, while 40% PEG 400 caused the appearance of a small minimum at 232 nm and disappearance of all other spectrum characteristics These changes are consistent with soluble aggregate formation with increasing quantities of PEG 400 (Urry et al., 1970; Urry and Ji, 1968).

The addition of glycerol resulted in the appearance of a more defined negative peak at 222 nm and the shifting of the disordered peak at 202 nm to 208 nm **(Figure 2F)**, indicating increased helicity as seen with TFE (Greenfield, 2006). The presence of an isodichroic point – a position where all spectra intersect– near 212 nm suggests that transitioning from a predominately disordered state to a helical structure is a reversible two-state process. These results are consistent with a model wherein increasing amounts of glycerol promote the folding of MSL10^N^ into a largely helical conformation and further demonstrate that the disordered MSL10^N^ exhibits structural sensitivity to a variety of solution environments.

### Phosphomimicry does not result in folding of MSL10^N^

Some IDRs undergo folding events in response to changes in phosphorylation state in order to regulate protein interaction (Bah et al., 2015). As phosphorylation of the N-terminus leads to inactivation of the MSL10 cell death signaling pathways, we investigated whether such phosphorylation-dependent folding occurs for MSL10^N^. We first compared the average per residue disorder scores of wild-type MSL10^N^, phosphodead (MSL10^N,7A^), and phosphomimetic (MSL10^N,7D^) sequences from 5 separate disorder predictors **(Figure 3A)**. Three out of five predictors showed that phosphomimetic MSL10^N,7D^ substitutions results in an approximately 0.75%-2.5% increase in propensity for disorder, while overall phosphodead MSL10^N,7A^ substitutions tended to decrease disordered propensity by 0.65-3.7% as compared to wild-type MSL10 ^N^. To determine whether phosphorylation affects the structure of the MSL10 N-terminus experimentally, we performed CD on purified His-tagged MSL10^N^, MSL10^N,7A^, and MSL10^N,7D^. During SDS-PAGE, MSL10^N^, MSL10^N,7A^, and MSL10^N,7D^ ran at exaggeratedly different speeds not predicted by their differences in molecular weight **(Figure 3B)**, which may be due to changes in the net negative charge of the protein from substituting phospho-sites. The CD spectra recorded for phospho-variants were highly similar to spectra recorded for wild-type MSL10^N^, implying that the N-terminus is largely disordered regardless of phosphorylation state **(Figure 3C)**.

**Figure 3.**
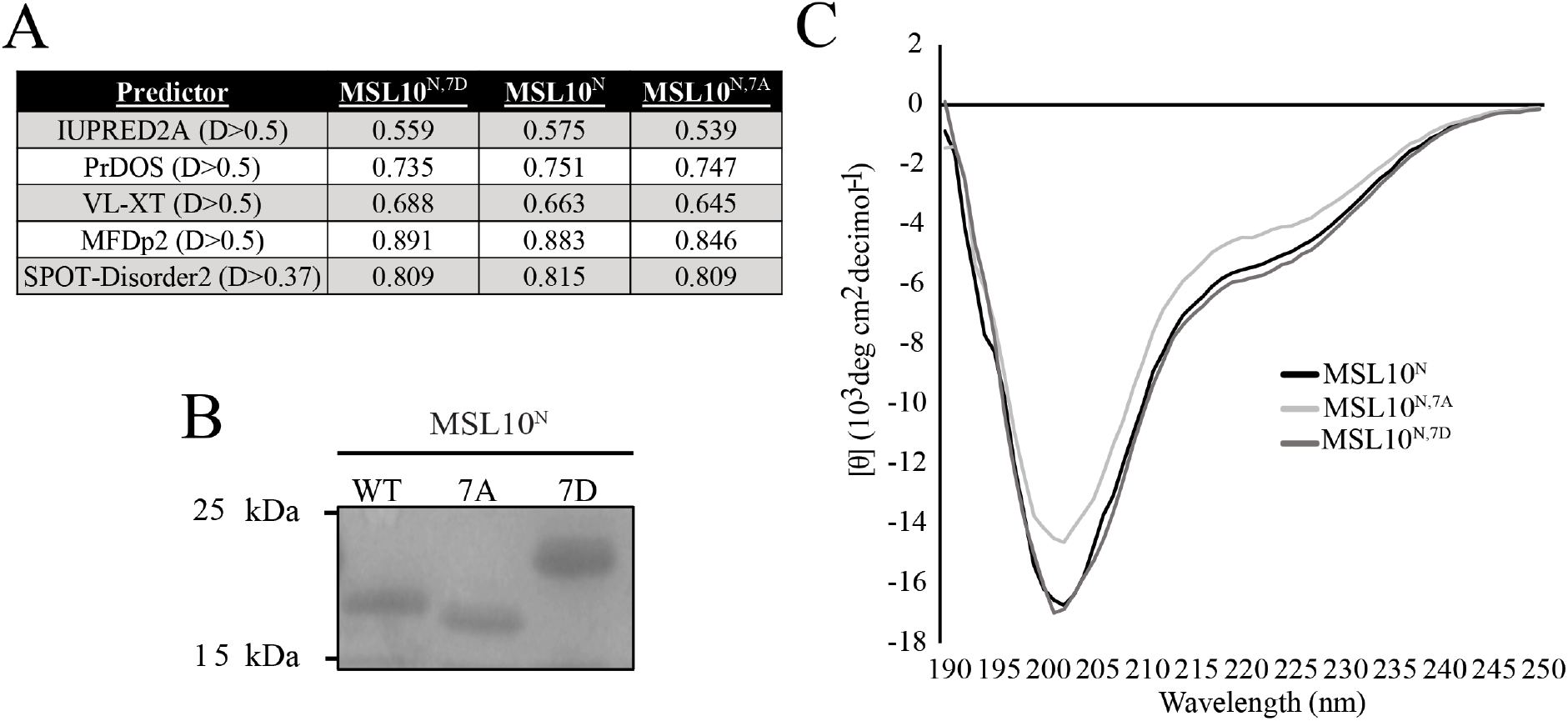
Phosphorylation does not result in folding of MSL10^N^. (A) Individual disorder scores for each amino acid in MSL10^N^were averaged for wild-type, phosphomimetic, and phosphodead variants using 5 prediction algorithms. Binary disorder-order prediction cutoffs for each predictor are noted in column headers, with scores larger than the given cutoff indicating disorder. (B) 10% Coomassie stained SDS-PAGE gel for His-tagged MSL10^N^, MSL10^N, 7D^, and MSL10^N, 7A^. (C) Circular dichroism spectra of His-tagged MSL10^N^, MSL10^N, 7D^, and MSL10^N, 7A^. Spectra were obtained at 20°C in 20 mM sodium phosphate buffer, pH 7.4.

### Aggregation and phase separation behavior of MSL8^N^, MSL9^N^, and MSL10^N^ *in vitro*

Some IDRs support networks of weak, multivalent interactions that result in condensate formation (Feng et al., 2019). Condensate formation has been implicated in a range of plant stress responses such as heat, salt, and water stress response (Emenecker et al., 2020; Cuevas-Velazquez and Dinneny, 2018; Dorone et al., 2021). Although poorly conserved on a sequence level, MSL8^N^, MSL9^N^, and MSL10^N^ exhibit similarities in sequence characteristics such as charge segregation, hydrophobicity, and the notable patterning of hydrophobic (aromatic) charged blocks along the sequence **(Figure 4A, Figure S1A)**. Such sequence characteristics are often found in IDRs that form condensates (Emenecker et al., 2020). Due to these sequence biases and to the observed ability of MSL10^N^ to aggregate via homotypic interaction following several different treatments in our CD assays, we probed MSL8^N^, MSL9^N^, and MSL10^N^ for their ability to form condensates *in vitro*.

**Figure 4.**
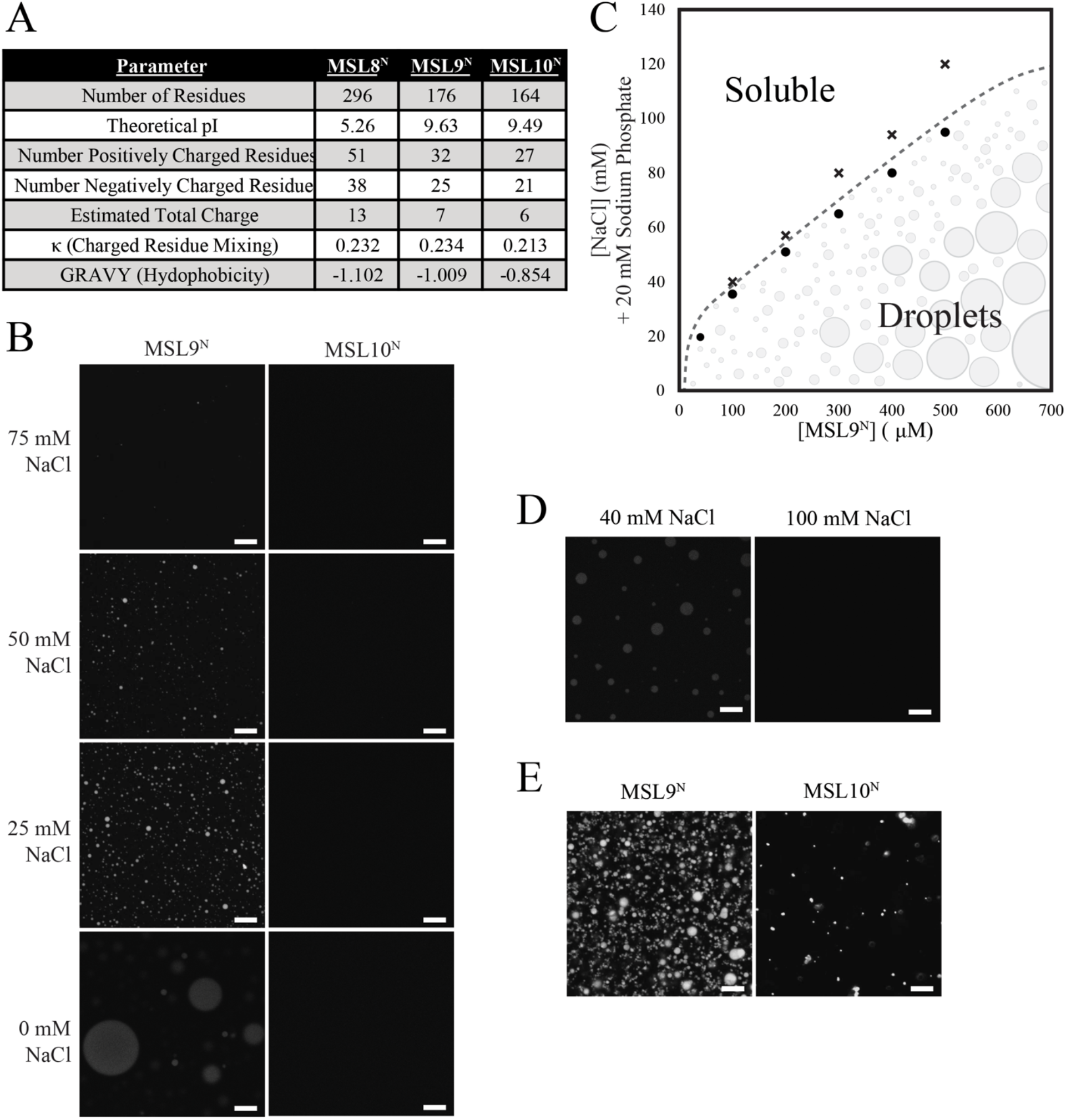
MSL9^N^ condensates in response to PEG 400 and low concentrations of NaCl, while MSL10^N^only forms assemblies with PEG 400 treatment. (A) Key parameters of MSL8^N^, MSL9^N^, and MSL10^N^ protein sequences as calculated by the ProtParam webtool. (B) Flourescence images of C-terminally His-tagged MSL9^N^ and MSL10^N^in different salt concentrations at ∼500 μM protein. (C) Phase diagram of MSL9^N^ as a function of protein and NaCl concentration. Black crosses denote tested conditions at which condensates were absent, while black circles are conditions that produced condensates. (D) Fluorescence images of MSL9^N^ wherein protein was treated with additional salt after droplet formation. Protein concentration before and after salt addition was ∼ 550 μ M. (E) Images of His-tagged MSL9^N^ and MSL10^N^ in response to 30% PEG 400 treatment at ∼ 500 μM protein. In B, D, and E, signal was detected at 500-540 nm, scale bars are 10 μm.

We first imaged purified proteins with fluorescence microscopy at various protein (42-1,000 μM) and NaCl (0-200 mM) concentrations as described in (Alberti et al., 2018). MSL10^N^ did not form any higher order assemblies in 0-200 mM NaCl and 500 μM protein **(Figure 4B, right panels)**. However, MSL9^N^ condensates formed in low NaCl concentrations at 500 μM protein **(Figure 4B, left panels)**. Condensates ranged in size, but were primarily ∼1 μM-20 μM in diameter. A MSL9^N^ phase diagram that further details the conditions at which MSL9^N^ forms condensates is depicted with an approximate phase boundary in **(Figure 4C)**. For some condensates, liquid-liquid phase separation (LLPS) – the spontaneous demixing of a homogenous solution into several phases that consist of different concentrations of a given protein component - drives formation. To determine if LLPS was the mechanism driving MSL9^N^ condensates, we assessed the reversibility of condensate formation – a property specifically exhibited by assemblies formed through LLPS. We first placed MSL9^N^ (550 μM) in 40 mM NaCl to induce the formation of condensates before adding MSL9^N^ stored in high-salt conditions to achieve a final concentration of 100 mM NaCl (with no concurrent change in MSL9^N^ concentration). Upon exposure to the 100 mM NaCl treatment, preformed MSL9^N^ condensates dispersed spontaneously, indicating that MSL9^N^ condensates form and disperse reversibly **(Figure 4D)**. This, in combination with the spherical nature of these condensates, implies that the MSL9^N^ condensates were liquid-like in character.

Due to our observations of MSL10^N^ aggregation in response to PEG 400 (**Figure 2E**), we assessed condensate formation following PEG treatment. In the presence of 30% PEG 400, MSL10^N^ formed clustered higher order assemblies ∼1-3 μM in diameter **(Figure 4E)**, further implying that increased molecular crowing may drive MSL10^N^ assembly. Notably, the addition of PEG 400 to MSL9^N^ samples resulted in spherical condensates of lesser volume (∼1-5 μM in diameter) than those seen in phosphate buffer alone at the same protein concentration. This implies that greater molecular crowding increases the critical concentration of protein required for MSL9^N^ condensate formation.

During sample preparation, MSL8^N^ was retained in inclusion bodies in the pellet when purified using the same protocol used for MSL9^N^ and MSL10^N^ **(Figure 5A, left panel)**. Therefore, we performed an insoluble preparation on the inclusion body-containing pellet fraction **(Figure 5A, right panel)**. A protein band around 99 kDa, which is exactly three times larger than a single MSL8^N^ protein, was consistently found across all elutions. In all salt conditions tested, MSL8^N^ formed non-spherical clustered assemblies **(Figure 5B)**, despite being at a lower protein concentration in our assays due to insoluble preparation. While both MSL8^N^ and MSL9^N^ showed condensate formation at low salt concentrations (25 mM NaCl), MSL8^N^ had a high propensity to form condensates even at low protein concentrations compared to MSL9^N^. We also noted that cold temperatures (4°C) caused the protein in storage solution to became turbid (**Figure 5C**). We therefore assessed the effect of temperature on MSL8^N^ condensate formation, with MSL9^N^ for comparison **(Figure 5D)**. We observed larger clustered assemblies in MSL8^N^ in cold temperatures compared to room temperature, indicating that MSL8^N^ condensate formation is temperature-dependent. Meanwhile, MSL9^N^ appeared slightly more soluble when exposed to cold temperatures. Collectively, our findings highlight that, even though they are all IDRs from the same domain of homologous proteins, MSL8^N^, MSL9^N^, and MSL10^N^ form condensates in response to strikingly different environmental conditions.

**Figure 5.**
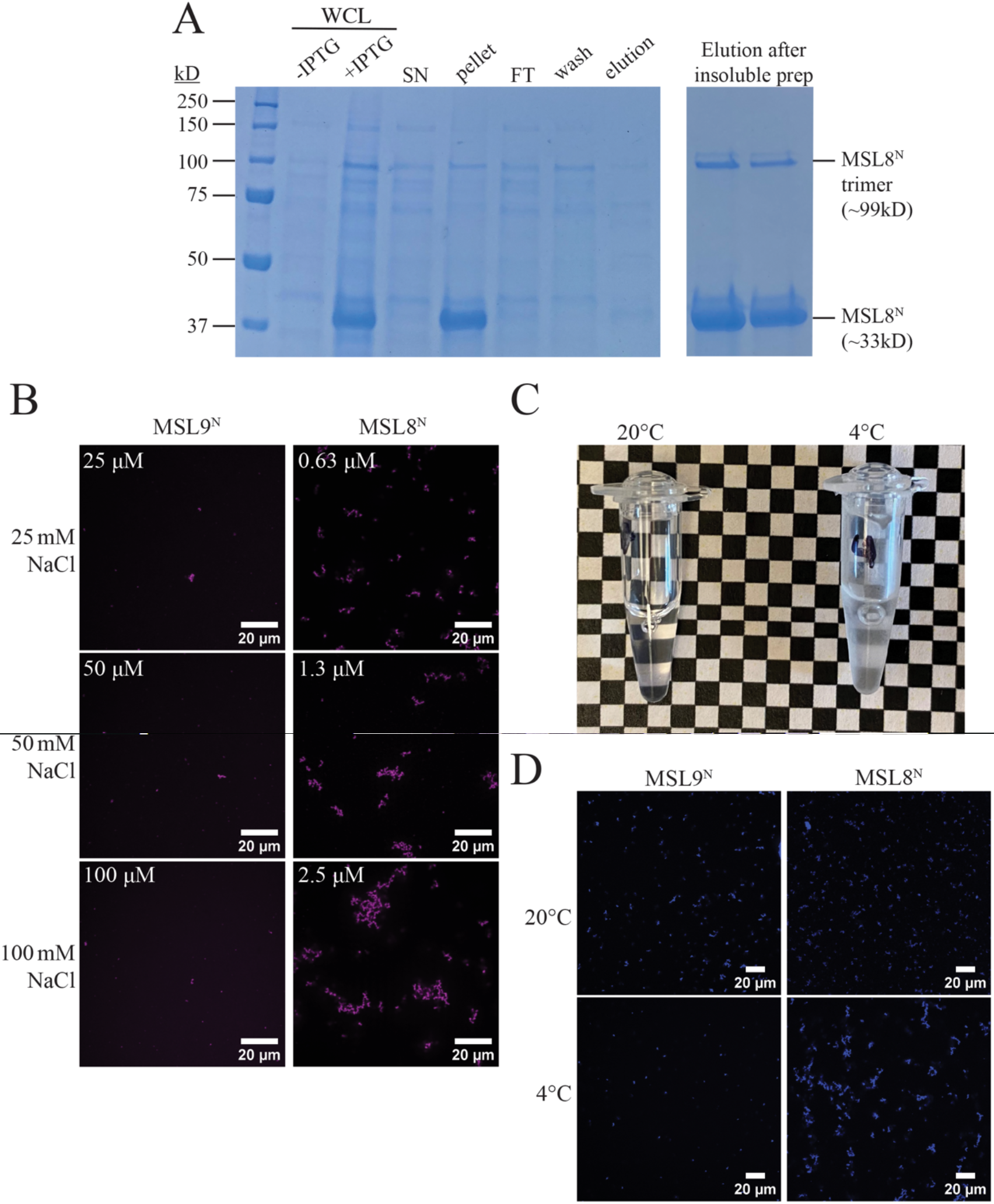
MSL8^N^ forms condensates in response to salt and to low temperatures. (A) Coomassie gel of samples taken during two MSL8^N^ purifications; soluble (left) and insoluble (right). WCL,whole cell lysate; SN, supematent; FT, flow-through. (B) Image of tubes containing purified MSL8^N^. The tube on the left was left at room temperature while the tube on the right was placed at 4°C. (C) Fluorescence images of His-tagged MSL9^N^ and MSL8^N^in increasing levels of salt in 20 mM sodium phosphate. Protein concentration is in upper left of each image. (D) Fluorescence images of MSL9^N^ and MSL8^N^ at room temperature (upper) and 4°C (lower). The samples in these images are in 50 mM NaCl, 20 mM sodium phosphate. The protein concentrations of MSL9^N^ and MSL8^N^ preparations were approximately 100 μM and 2.5 μM, respectively.

## DISCUSSION

### The soluble N-termini of MSL8, 9, and 10 are environmentally sensitive intrinsically disordered regions

Although the study of intrinsic disorder in membrane proteins such as ion channels has grown in the last several years, this area of research remains underappreciated. Here, we document the presence and characterize the properties of IDRs in the soluble N-terminus of *Arabidopsis thaliana* mechanosensitive ion channels, adding to our understanding of IDR properties found at the membrane. We consistently predicted intrinsic disorder in the soluble N-termini of MSL8, MSL9, and MSL10 – as well as MSL orthologs throughout the plant lineage **(Figure 1)**.

Although the primary sequence of MSL N-termini is poorly conserved both within a given species (Arabidopsis) and across plants **(Figure S1)**, disorder appears to be a trait that was maintained throughout evolutionary history. In contrast to structured regions, disordered sequences are typically functionally driven by bulk sequence features such as the amino acid composition, charge, or presence of disorder in the region; thus, their primary sequence can experience large perturbations without effect on IDR function (Zarin et al., 2017). Therefore, we predicted that disorder, but not primary sequence, is a conserved feature of the N-termini of MSLs.

These predictions were born out in circular dichroism experiments with purified recombinant protein of the exemplar MSL10^N^ **(Figure 2)**. MSL10^N^ spectra exhibited a broad negative peak that is indicative of disorder as the main form of secondary structure in the chain. The presence of a slight shoulder in the spectra indicated some helical content in the chain, but whether this helix is transiently formed or stable – as well as where helixes occur in the chain – requires further study.

Additionally, MSL10^N^ folds in the presence of the small molecule glycerol **(Figure 2F)**. Responses to crowding and osmolyte changes *in vitro* have been documented in other intrinsically disordered plant proteins, most notably in the Late Embryogenesis Abundant (LEA) and FLOE1 families (Rivera-Najera et al., 2014; Artur et al., 2019; Koubaa et al., 2019; Dorone et al., 2021). Some instances of environmentally driven folding are also seen in the disordered ASR proteins associated with abscisic acid signaling during fruit ripening and drought response (Hamdi et al., 2017). MSL10^N^ provides another example of an intrinsically disordered region from plants that is susceptible to conformational changes in response to physiochemical environmental modulations *in vitro*. Future studies will reveal if these traits are relevant *in vivo*.

### Relevance of N-terminal disorder in mediating transient protein interactions

Molecular Recognition Features (MoRFs), small 5-25 residue sequences that transition from a disordered to ordered state upon interacting with a binding partner, are regions found almost exclusively within IDRs (Mohan et al., 2006; Yang et al., 2019) and are important in some binding events and IDR-mediated protein complex formation (Mohan et al., 2006). The helical folding of MSL10^N^ in response to TFE **(Figure 2D)** suggests that the N-terminus could participate in such disorder-to-order-inducing interactions. The disordered binding region predictor ANCHOR2 (Mészáros et al., 2018) predicted that the first 60 residues of MSL10^N^ and residues 70-95 comprise protein binding regions and MoRFPred and MoRFChibi–Web tools (Disfani et al., 2012; Malhis et al., 2016) predicted MoRFs at amino acids 1-33, 74-91, and 134-145 **(Figure S4)**. MoRFs are predicted in the same regions of all MSL9 and MSL10 orthologs investigated, suggesting that protein interaction through MoRFs may be conserved (**Figure S4B**).

Conserved motifs within IDRs often serve as modules of protein interaction (Ahrens et al., 2017). The conserved putative MoRF regions in MSL10^N^ and its orthologs coincided with the most conserved regions of the otherwise poorly conserved N-termini, including amino acids 15-24, 60-65, and 72-92 **(Figure S1B, Figure S4B)**. In addition, all three predicted MoRF regions in Arabidopsis MSL10^N^ either contained or were adjacent to at least one N-terminal phosphorylation site previously identified by proteomic analyses (S29, S57, S128, S131, T136) (summarized in (Veley et al., 2014)) **(Figure 3)**. This is of interest as phosphorylation may act as a key regulatory factor in MoRF transitions and function (Mohan et al., 2006; Wright and Dyson, 2015; Darling and Uversky, 2018).

### The N-termini of MSL8, MSL9 and MSL10 show distinct assembly characteristics

Unexpectedly, His-tagged MSL8^N^, MSL9^N^, and MSL10^N^ undergo condensate formation in sharply different physiochemical environments *in vitro* **(Figure 4, Figure 5)**. While MSL10^N^ did not respond to changes in salt concentrations in the tested conditions, it did form assemblies when introduced to environments with higher molecular crowding in both CD and *in vitro* imaging **(Figure 2E, Figure 4E)**. Additionally, our CD data **(Figure 2B)** imply that MSL10^N^ may form condensates with increasing temperature. Conversely, MSL9^N^ spontaneously formed condensates in low salt conditions, likely via LLPS **(Figure 4B-D)**. The addition of PEG 400 decreased condensate size at a given concentration of salt, implying that crowding reduces the propensity of MSL9^N^ to undergo LLPS **(Figure 4E)**. However, molecular crowding agents such as PEG typically have pleiotropic effects that may be responsible for the observed behavior (Zhou et al., 2008; Christiansen et al., 2013). MSL9^N^ also showed decreased condensate formation at low temperatures **(Figure 5D)**. Lastly, MSL8^N^ required insoluble preparation for purification and formed clustered higher order assemblies when presented with low salt, low protein, and low temperature conditions **(Figure 5)**. To summarize, all three N-terminal domains tested here exhibited distinct characteristics of condensate formation.

As noted above, MSL8^N^, MSL9^N^, and MSL10^N^ show little sequence conservation, despite evolutionary relatedness and conserved characteristics such as intrinsic disorder, hydrophobicity, and charge segregation **(Figure 4A, Figure S1A)**. These sequence differences may drive how each of the IDRs responds distinctively to various environmental contexts. For example, and in line with the stickers and spacers model of phase separation (Emenecker et al., 2020), several charged and aromatic residues found in MSL9^N^ (R4, F31, Y60, F62, R80, F102, Y104, R108, R114, F120, F125, R127) but absent in MSL10^N^ may be responsible for discrepancies in assembly between the two MSL family members. Additionally, insertions in MSL8^N^ relative to MSL9^N^ and MSL10^N^ have contributed a series of additional aromatic and charged sites that may confer a greater propensity to form condensates **(Figure S1A)**. Future work should target these residues as potential sequence determinants of phase separation.

### Potential relevance of MSL^N^ assembly *in vivo*

Phase separation at the membrane can serve several biological functions. Condensate formation may be regulated by or promote certain protein interactions or biochemical reactions such as post translational modification (Alberti and Hyman, 2021). Alternatively, phase separation at the membrane sometimes contributes to membrane bending (Yuan et al., 2021; Feng et al., 2019).

Future studies will determine if such IDR molecular functions of are relevant to MSL10’s physiological function during cell swelling in seedlings (Basu and Haswell, 2020), MSL8’s function during pollen grains hydration (Hamilton et al., 2015), or to the unknown functions of MSL9.

The findings shown here provide insight into the structural character of the N-termini of three members of the Arabidopsis MSL family of mechanosensitive ion channels. Each of the three channels studied here have different expression patterns in the plant and are involved in distinct functions within those tissues; they also have very different N-terminal IDR sequences that exhibit distinct behaviors. This study sets the stage for research into the relationship between *in vitro* MSL IDR behavior and *in vivo* function.

## METHODS

### Computational analyses

Predictions of intrinsic disorder were performed using several webserver-based predictors, including IUPred2A (https://iupred2a.elte.hu/), FoldIndex (https://fold.weizmann.ac.il/fldbin/findex), PONDR-VLXT (http://www.pondr.com/), SPOT-Disorder2 (https://sparks-lab.org/server/spot-disorder2/), PrDOS (http://prdos.hgc.jp/cgi-bin/top.cgi), CSpritz (http://old.protein.bio.unipd.it/cspritz/), PONDR-FIT (http://original.disprot.org/pondr-fit.php), DISOPRED3 (http://bioinf.cs.ucl.ac.uk/psipred/), and MFDp2 (http://biomine.cs.vcu.edu/servers/MFDp2/). Protein binding region and MoRF predictions were made using the ANCHOR2 (https://iupred2a.elte.hu/), MoRFPred (http://biomine.cs.vcu.edu/servers/MoRFpred/), and MoRFChibi_Web (https://morf.msl.ubc.ca/index.xhtml) algorithms. The PredictProtein (https://predictprotein.org/) and Composition Profiler (http://www.cprofiler.org/cgi-bin/profiler.cgi) webservers were used for analysis of the amino acid composition of the MSL N-termini. Topology predictions were performed using the TMHMM 2.0 (http://www.cbs.dtu.dk/services/TMHMM/) and ARAMEMNON webservers (http://aramemnon.uni-koeln.de/). The CIDER (http://pappulab.wustl.edu/CIDER/) webserver was used for Das-Pappu plot generation and calculation of κ-values (Holehouse et al., 2017). To compare the global propensities of disorder of the wild-type, phosphodead, and phosphomimetic variants of the MSL10 N-terminus, disorder scores for all residues of each N-terminal variant (residues 1-164) were averaged and reported for 5 different prediction algorithms.

### Alignments and aligned disorder scoring

MSL8, MSL9, and MSL10 sequences were aligned using the blastp algorithm in the BLAST webserver (Altschul et al., 1990); one position was manually corrected (gap shifted at aligned positions 144-149). The ClustalW program embedded within the MEGA7 software package was used to align *Arabidopsis thaliana* MSL10 and putative plant orthologs reported in (Basu et al., 2020), (Pairwise alignment – Gap Opening Penalty = 15 and Gap Extension Penalty = 0.5; Multiple alignment – Gap Opening Penalty = 15 and Gap Extension Penalty = 0.2; Residue-specific penalties and Hydrophilic penalties ON; Gap separation difference = 4). To plot the disorder scores of MSL8 and MSL9 and to find the median IUPRED2A disorder score of all MSL9 and MSL10 orthologs, the process described in (Christensen et al., 2019) was applied. Briefly, at an aligned position in *Arabidopsis thaliana* MSL10, the alignment of *Arabidopsis thaliana* MSL10 with each individual ortholog was manually compared, and non-aligning regions were removed from consideration. The disorder scores for all ortholog residues that aligned with a position of MSL10 were medialized and reported for that aligned position.

### MSL9^N^ and MSL10^N^ protein expression and purification

*E. coli* strain DE3 (Rosetta) chemically competent cells were transformed with pET-26b(+) plasmids containing the protein domain of interest. Overnight cultures were diluted into sterilized LB media and incubated with shaking at 37°C until reaching an optical density at 600 nm (OD_600_) of approximately 0.5. 1 mM isopropyl β-d-thiogalactopyranoside (IPTG) was added to the grown cultures, and protein induction was carried out at 37°C with shaking for 2 hours. Cultured cells were flash frozen prior to storage at -80°C.

For purification, each pellet was thawed over ice and resuspended in 50 mL of lysis buffer (50 mM sodium phosphate, 1 mg/mL lysozyme, 0.25% Tween 20, 300 mM NaCl, and 10 mM imidazole; supplemented with 0.1 mM PMSF, 3 µM Leupeptin, and 1 µM Pepstatin). Resuspended pellets were incubated on ice for 30 minutes and then sonicated 5-10 times (1 second pulses, 50% amplitude, 10 second timer) until pale yellow in color, and debris was spun down. The resulting supernatant was added to tubes containing HisPur NiNTA resin (Thermo Fisher Scientific) and tubes were rocked for 30 minutes at 4°C. Supernatant was removed and resin was washed three times. Protein was eluted with elution buffer (50 mM sodium phosphate, 300 mM NaCl, 250 mM imidazole). Eluates were stored at 4°C overnight before further sample preparation steps took place. For eluates placed in long term storage, 50% glycerol was added to eluates to achieve a final sample containing 10% glycerol before flash freezing in liquid nitrogen and storing at -80°C.

### Far-UV circular dichroism sample preparation and measurements

Ni-NTA eluates were thawed on ice and desalted via the PD-10 desalting column system (GE Healthcare) following the manufacturer’s gravity flow protocol. Following initial desalting, samples were subjected to Pierce PES 2-6 mL concentrator columns (Thermo Fisher Scientific) to further desalt the sample, concentrate the protein to 1-2 mg/mL, and buffer swap into 20 mM sodium phosphate buffer (pH = 7.4). Final sample purities and concentrations were estimated with Coomassie staining of SDS-PAGE gels, Bradford assays, and UV absorbance measurements at 280 nm performed in triplicate on a Nanodrop One Microvolume UV-Vis Spectrophotometer (Thermo Fisher Scientific).

CD was performed using the JASCO J-810 CD Spectrometer set at 20°C. A 0.1 cm path length cuvette was measured over the range of 190 nm – 260 nm. His-tagged protein samples were used in the concentration range of 0.1 mg/mL – 0.3 mg/mL protein, as estimated by Bradford Assay. Six accumulations were taken and averaged for each measurement. Spectra were deconvoluted using the DichroWeb tool, using three separate algorithms (CONTIN, SELCON3, CDSSTR) for each spectrum to confirm analysis consistency; calculations were performed using reference set 7. For variable temperature circular dichroism, a single aliquot of purified MSL10 N-terminus was exposed to progressively increasing temperatures, from 10°C to 80°C. For conditional CD experiments (including TFE, PEG 400, and glycerol treatments), measurements made at each percentage of a given treatment were conducted with a separate aliquot of purified protein. As proposed by (Bruch et al., 1991), the ratio of values found at 208 nm and 222 nm was calculated to assess the character of alpha-helical content present within the analyzed protein.

### Sample preparation and microscopy for *in vitro* condensate imaging for MSL9 and MSL10

Size exclusion chromatography was performed on a HiLoad 16/600 Superdex 200 pg (GE Life Sciences) connected to an AKTA FPLC (GE Life Sciences) to further purify samples isolated via His-tag purification and buffer swap into a “high salt” buffer (200 mM NaCl, 20 mM Sodium Phosphate, pH = 7.5). The concentrations and purities of fractions were assessed via absorbance at 280 nm and Coomassie staining, respectively, before pooling desired fractions and concentrating with Pierce PES 2-6 mL concentrator columns (Thermo Fisher Scientific). Concentrated protein was aliquoted and flash frozen in liquid N_2_ before storage at -80°C.

For imaging, His-tagged protein was labeled with Alexa Fluor 488 NHS Ester (Thermo Fisher Scientific) per manufacturer’s recommendations. Labeled and unlabeled variants of a given His-tagged protein were mixed in a molar ratio of 1:400. In a 1.6 mm deep silicon isolator well (Grace Bio-Labs) adhered to a standard microscope slide, protein in 200 mM NaCl, 20 mM sodium phosphate, pH = 7.5 was diluted with 20 mM sodium phosphate (with some concentration of NaCl, if required) to achieve the desired combination of NaCl and protein concentration. Similarly, samples prepared in 20 mM sodium phosphate were mixed with PEG 400 to achieve a final PEG concentration of 30%. Mounted samples were then imaged immediately using the 20X objective of an Olympus IX83 confocal microscope at room temperature. Image brightness and contrast were adjusted (equivalently for all images) in post to improve visibility of condensates in monochromatic color scheme.

### MSL8^N^ expression and purification

*E. coli* strain DE3 (Rosetta) chemically competent cells were transformed with pET-26b(+) plasmids containing the protein domain of interest (MSL8^N^ or MSL9^N^). Overnight cultures were diluted into sterilized LB media and incubated with shaking at 37°C until reaching an optical density at 600 nm (OD_600_) of approximately 0.5. 1 mM isopropyl β-d-thiogalactopyranoside (IPTG) was added to the grown cultures, and protein induction was carried out at 37°C with shaking for 2 hours. Cultured cells were flash frozen prior to storage at -80°C. For purification, each pellet was thawed over ice and resuspended in 50 mL of lysis buffer (50 mM sodium phosphate, 1 mg/mL lysozyme, 0.25% Tween 20, 300 mM NaCl, and 10 mM imidazole; supplemented with 0.1 mM PMSF, 3 µM Leupeptin, and 1 µM Pepstatin). Resuspended pellets were incubated on ice for 30 minutes and then sonicated 5-10 times (1 second pulses, 50% amplitude, 10 second timer) until pale yellow in color, and the remaining pellet was spun down.

For MSL8^N^, an insoluble preparation was performed on this pellet (see below). For MSL9^N^, the resulting supernatant was added to tubes containing HisPur NiNTA resin (Thermo Fisher Scientific) and tubes were rocked for 30 minutes at 4°C. Supernatant was removed and resin was washed three times. Protein was eluted with elution buffer (50 mM sodium phosphate, 300 mM NaCl, 250 mM imidazole). The pellet containing MSL8^N^ inclusion bodies was solubilized into a resuspension buffer containing 6 M GdmCl, 20 mM Tris, and 15 mM imidazole (pH 7.5) using sonication (3 times, 1 second on and 2 seconds off, 30% amplitude, 1 min timer). The resulting suspension was then clarified via centrifugation (35,000 x g for 45 min). The lysate was applied to HisPur NiNTA resin (Thermo Fisher; Cat#88221) in a gravity column for 10 min, the solution was then collected and run through again before saving as “flow through”. The column was washed with buffer containing 20 mM Tris, 30 mM imidazole, and 4 M urea (pH 7.5) until protein was no longer detected via blue coloration in Bradford reagent droplets. Protein was eluted using an elution buffer containing 20 mM Tris, 350 mM imidazole, 4 M urea (pH 7.5). All eluates were stored at 4°C overnight before further sample preparation steps took place. For eluates placed in long term storage, 50% glycerol was added to eluates to achieve a final sample containing 10% glycerol before flash freezing in liquid nitrogen and storing at -80°C. The purities of fractions were assessed via Coomassie staining before pooling desired fractions and concentrating with 0.5 mL Peirce PES concentrator columns (Thermo Fisher; Cat#88513). For unlabeled protein, samples were buffer swapped into 200 mM NaCl, 20 mM Sodium Phosphate using 0.5 mL Zeba spin desalting columns (Thermo Fisher; Cat#89882) immediately before imaging.

### MSL8^N^ *in vitro* condensate imaging

For labeled protein samples, both MSL8^N^ and MSL9^N^ were placed in a high salt storage buffer (1 M NaCl, 200 mM Sodium Phosphate) using gravity PD-10 desalting columns (GE Healthcare). The protein was labeled with Alexa Fluor 647 NHS (Invitrogen; Cat#A37573) per manufacturer’s recommendations. Any MSL8^N^ condensates that formed in the cold during labeling were spun down and removed. Labeled protein was aliquoted before flash-freezing and storing at -80°C. Labeled and unlabeled variants of a given His-tagged protein were mixed in a molar ratio of 1:400. In a 1.7 mm deep silicon isolator well (Grace Bio-Labs; SKU#665201) adhered to a standard microscope slide, protein in 200 mM NaCl, 20 mM sodium phosphate, pH = 7.5 was diluted with 20 mM sodium phosphate to achieve the desired combination of NaCl and protein concentration. Mounted samples were then imaged immediately using the 20X objective of a Nikon T2i microscope confocal microscope at room temperature or with an added ice pack to cool the sample to 4°C. Image brightness and contrast were adjusted (equivalently for all images) to improve visibility of condensates.

### GenBank accessions

The sequences for MSL8 (At2G17010.1), MSL9 (AT5G19520.1), and MSL10 (AT5G12080.1) can be found on The Arabidopsis Information Resource (TAIR). MSL9 and MSL10 orthologous sequences used for predictions and alignments can be found at the following GenBank accession numbers: *Arabidopsis thaliana –* NP_196769.1; *Arabidopsis lyrata –* XP_002873549.1; *Brassica rapa –* XP_009121883.1; *Brassica napus* – XP_013676093.1; *Camelina sativa* – XP_010453270.1; *Medicago truncatula* – XP_024633438.1; *Vitis vinifera* – XP_002279755.1; *Solanum lycopersicum* – XP_004245056.1; *Solanum tuberosum* XP_006350354.1; *Zea mays* – XP_008649202.1; *Oryza sativa –* XP_015641284.1; *Sorghum bicolor* – XP_002438025.1; *Setaria italica* – XP_004964936.1; *Brachypodium distachyon* – XP_003560953.1.

## ACKNOWLEDGMENTS

This work was supported by NSF Graduate Research Fellowship (DGE-2139839 and DGE-1745038) and the William H. Danforth Fellowship in Plant Sciences to K.M., NSF MCB 1929355 to E.S.H., and the NSF Center for Engineering Mechanobiology (NSF grant CMMI 1548571). We would like to thank Ryan Emenecker for fantastic scientific discussions during the course of this work.

## Notes

### Competing Interest Statement

The authors have declared no competing interest.

## REFERENCES

Ahrens, J.B., Nunez-Castilla, J., and Siltberg-Liberles, J. (2017). Evolution of intrinsic disorder in eukaryotic proteins. Cell Mol Life Sci 74: 3163–3174.

Alberti, S. and Hyman, A.A. (2021). Biomolecular condensates at the nexus of cellular stress, protein aggregation disease and ageing. Nat Rev Mol Cell Bio 22: 196–213.

Alberti, S., Saha, S., Woodruff, J.B., Franzmann, T.M., Wang, J., and Hyman, A.A. (2018). A User’s Guide for Phase Separation Assays with Purified Proteins. J Mol Biol 430: 4806–4820.

Altschul, S.F., Gish, W., Miller, W., Myers, E.W., and Lipman, D.J. (1990). Basic local alignment search tool. J Mol Biol 215: 403–410.

Artur, M.A.S., Rienstra, J., Dennis, T.J., Farrant, J.M., Ligterink, W., and Hilhorst, H. (2019). Structural Plasticity of Intrinsically Disordered LEA Proteins from Xerophyta schlechteri Provides Protection In Vitro and In Vivo. Front Plant Sci 10: 1272.

Bah, A., Vernon, R.M., Siddiqui, Z., Krzeminski, M., Muhandiram, R., Zhao, C., Sonenberg, N., Kay, L.E., and Forman-Kay, J.D. (2015). Folding of an intrinsically disordered protein by phosphorylation as a regulatory switch. Nature 519: 106–109.

Basu, D., Codjoe, J.M., Veley, K.M., and Haswell, E.S. (2022). The Mechanosensitive Ion Channel MSL10 Modulates Susceptibility to Pseudomonas syringae in Arabidopsis thaliana. Mol Plant-microbe Interactions 35: 567–582.

Basu, D. and Haswell, E.S. (2017). Plant mechanosensitive ion channels: an ocean of possibilities. Current Opinion in Plant Biology 40: 43–48.

Basu, D. and Haswell, E.S. (2020). The Mechanosensitive Ion Channel MSL10 Potentiates Responses to Cell Swelling in Arabidopsis Seedlings. Current Biology 30: 2716-2728.e6.

Basu, D., Shoots, J.M., and Haswell, E.S. (2020). Interactions between the N- and C-termini of mechanosensitive ion channel AtMSL10 are consistent with a three-step mechanism for activation. Journal of Experimental Botany.

Bruch, M.D., Dhingra, M.M., and Gierasch, L.M. (1991). Side chain–backbone hydrogen bonding contributes to helix stability in peptides derived from an α-helical region of carboxypeptidase A. Proteins Struct Funct Bioinform 10: 130–139.

Bürgi, J., Xue, B., Uversky, V.N., and Goot, F.G. van der (2016). Intrinsic Disorder in Transmembrane Proteins: Roles in Signaling and Topology Prediction. Plos One 11: e0158594.

Chemes, L.B., Alonso, L.G., Noval, M.G., and Prat-Gay, G. de (2012). Circular dichroism techniques for the analysis of intrinsically disordered proteins and domains. Methods Mol Biology Clifton N J 895: 387–404.

Christensen, L.F., Staby, L., Bugge, K., O’Shea, C., Kragelund, B.B., and Skriver, K. (2019). Evolutionary conservation of the intrinsic disorder-based Radical-Induced Cell Death1 hub interactome. Sci Rep-uk 9: 18927.

Christiansen, A., Wang, Q., Cheung, M.S., and Wittung-Stafshede, P. (2013). Effects of macromolecular crowding agents on protein folding in vitro and in silico. Biophysical Rev 5: 137–145.

Cohan, M.C., Ruff, K.M., and Pappu, R.V. (2019). Information theoretic measures for quantifying sequence–ensemble relationships of intrinsically disordered proteins. Protein Eng Des Sel 32: 191–202.

Covarrubias, A.A., Cuevas-Velazquez, C.L., Romero-Pérez, P.S., Rendón-Luna, D.F., and Chater, C.C.C. (2017). Structural disorder in plant proteins: where plasticity meets sessility. Cell Mol Life Sci 74: 3119–3147.

Cuevas-Velazquez, C.L. et al. (2021). Intrinsically disordered protein biosensor tracks the physical-chemical effects of osmotic stress on cells. Nat Commun 12: 5438.

Cuevas-Velazquez, C.L. and Dinneny, J.R. (2018). Organization out of disorder: liquid-liquid phase separation in plants. Current Opinion in Plant Biology 45: 68–74.

Cuevas-Velazquez, C.L., Saab-Rincón, G., Reyes, J.L., and Covarrubias, A.A. (2016). The Unstructured N-terminal Region of Arabidopsis Group 4 Late Embryogenesis Abundant (LEA) Proteins Is Required for Folding and for Chaperone-like Activity under Water Deficit*. J Biol Chem 291: 10893–10903.

Darling, A.L. and Uversky, V.N. (2018). Intrinsic Disorder and Posttranslational Modifications: The Darker Side of the Biological Dark Matter. Frontiers Genetics 9: 158.

Das, R.K. and Pappu, R.V. (2013). Conformations of intrinsically disordered proteins are influenced by linear sequence distributions of oppositely charged residues. Proc National Acad Sci 110: 13392–13397.

Das, R.K., Ruff, K.M., and Pappu, R.V. (2015). Relating sequence encoded information to form and function of intrinsically disordered proteins. Curr Opin Struc Biol 32: 102–112.

Disfani, F.M., Hsu, W.-L., Mizianty, M.J., Oldfield, C.J., Xue, B., Dunker, A.K., Uversky, V.N., and Kurgan, L. (2012). MoRFpred, a computational tool for sequence-based prediction and characterization of short disorder-to-order transitioning binding regions in proteins. Bioinformatics 28: i75–i83.

Dorone, Y. et al. (2021). A prion-like protein regulator of seed germination undergoes hydration-dependent phase separation. Cell 184: 4284-4298.e27.

Dunker, A.K. et al. (2013). What’s in a name? Why these proteins are intrinsically disordered: Why these proteins are intrinsically disordered. Intrinsically Disord Proteins 1: e24157.

Emenecker, R.J., Holehouse, A.S., and Strader, L.C. (2020). Emerging Roles for Phase Separation in Plants. Dev Cell 55: 69–83.

Feng, Z., Chen, X., Wu, X., and Zhang, M. (2019). Formation of biological condensates via phase separation: Characteristics, analytical methods, and physiological implications. J Biol Chem 294: 14823–14835.

Goretzki, B., Guhl, C., Tebbe, F., Harder, J.-M., and Hellmich, U.A. (2021). Unstructural biology of TRP ion channels: The role of intrinsically disordered regions in channel function and regulation. J Mol Biol: 166931.

Greenfield, N.J. (2006). Using circular dichroism spectra to estimate protein secondary structure. Nat Protoc 1: 2876–2890.

Hamdi, K., Salladini, E., O’Brien, D.P., Brier, S., Chenal, A., Yacoubi, I., and Longhi, S. (2017). Structural disorder and induced folding within two cereal, ABA stress and ripening (ASR) proteins. Sci Rep-uk 7: 15544.

Hamilton, E.S., Jensen, G.S., Maksaev, G., Katims, A., Sherp, A.M., and Haswell, E.S. (2015). Mechanosensitive channel MSL8 regulates osmotic forces during pollen hydration and germination. Science (New York, N.Y.) 350: 438–441.

Haswell, E.S. (2007). MscS-Like Proteins in Plants. In Mechanosensitive Ion Channels, Part A. (Elsevier), pp. 329–359.

Haswell, E.S. and Meyerowitz, E.M. (2006). MscS-like proteins control plastid size and shape in Arabidopsis thaliana. Current Biology 16: 1–11.

Haswell, E.S., Peyronnet, R., Barbier-Brygoo, H., Meyerowitz, E.M., and Frachisse, J.-M. (2008). Two MscS Homologs Provide Mechanosensitive Channel Activities in the Arabidopsis Root. Curr Biol 18: 730–734.

Holehouse, A.S., Das, R.K., Ahad, J.N., Richardson, M.O.G., and Pappu, R.V. (2017). CIDER: Resources to Analyze Sequence-Ensemble Relationships of Intrinsically Disordered Proteins. Biophys J 112: 16–21.

Hua, Q.-X., Jia, W., Bullock, B.P., Habener, J.F., and Weiss, M.A. (1998). Transcriptional Activator−Coactivator Recognition: Nascent Folding of a Kinase-Inducible Transactivation Domain Predicts Its Structure on Coactivator Binding †. Biochemistry-us 37: 5858–5866.

Kjaergaard, M. and Kragelund, B.B. (2017). Functions of intrinsic disorder in transmembrane proteins. Cell Mol Life Sci 74: 3205–3224.

Koubaa, S., Bremer, A., Hincha, D.K., and Brini, F. (2019). Structural properties and enzyme stabilization function of the intrinsically disordered LEA_4 protein TdLEA3 from wheat. Sci Rep-uk 9: 3720.

Lau, S.Y., Taneja, A.K., and Hodges, R.S. (1984). Synthesis of a model protein of defined secondary and quaternary structure. Effect of chain length on the stabilization and formation of two-stranded alpha-helical coiled-coils. J Biol Chem 259: 13253–13261.

Lee, C.P., Maksaev, G., Jensen, G.S., Murcha, M.W., Wilson, M.E., Fricker, M., Hell, R., Haswell, E.S., Millar, A.H., and Sweetlove, L.J. (2016). MSL1 is a mechanosensitive ion channel that dissipates mitochondrial membrane potential and maintains redox homeostasis in mitochondria during abiotic stress. The Plant Journal 88: 809–825.

Lee, R. van der et al. (2014). Classification of Intrinsically Disordered Regions and Proteins. Chem Rev 114: 6589–6631.

Luo, P. and Baldwin, R.L. (1997). Mechanism of Helix Induction by Trifluoroethanol: A Framework for Extrapolating the Helix-Forming Properties of Peptides from Trifluoroethanol/Water Mixtures Back to Water †. Biochemistry-us 36: 8413–8421.

Maksaev, G. and Haswell, E.S. (2012). MscS-Like10 is a stretch-activated ion channel from Arabidopsis thaliana with a preference for anions. Proceedings of the National Academy of Sciences of the United States of America 109: 19015–19020.

Maksaev, G., Shoots, J.M., Ohri, S., and Haswell, E.S. (2018). Nonpolar residues in the presumptive pore-lining helix of mechanosensitive channel MSL10 influence channel behavior and establish a nonconducting function. Plant Direct 2: 1–13.

Malhis, N., Jacobson, M., and Gsponer, J. (2016). MoRFchibi SYSTEM: software tools for the identification of MoRFs in protein sequences. Nucleic Acids Res 44: W488–W493.

Mansouri, A.L., Grese, L.N., Rowe, E.L., Pino, J.C., Chennubhotla, S.C., Ramanathan, A., O’Neill, H.M., Berthelier, V., and Stanley, C.B. (2016). Folding propensity of intrinsically disordered proteins by osmotic stress. Molecular bioSystems 12: 3695–3701.

Mao, A.H., Lyle, N., and Pappu, R.V. (2012). Describing sequence–ensemble relationships for intrinsically disordered proteins. Biochem J 449: 307–318.

Mészáros, B., Erdős, G., and Dosztányi, Z. (2018). IUPred2A: context-dependent prediction of protein disorder as a function of redox state and protein binding. Nucleic Acids Res 46: W329–W337.

Mohan, A., Oldfield, C.J., Radivojac, P., Vacic, V., Cortese, M.S., Dunker, A.K., and Uversky, V.N. (2006). Analysis of Molecular Recognition Features (MoRFs). J Mol Biol 362: 1043–1059.

Na, J.-H., Lee, W.-K., and Yu, Y. (2018). How Do We Study the Dynamic Structure of Unstructured Proteins: A Case Study on Nopp140 as an Example of a Large, Intrinsically Disordered Protein. International Journal of Molecular Sciences 19: 381–11.

Nakano, S., Miyoshi, D., and Sugimoto, N. (2014). Effects of Molecular Crowding on the Structures, Interactions, and Functions of Nucleic Acids. Chem Rev 114: 2733–2758.

Pivetti, C.D., Yen, M.-R., Miller, S., Busch, W., Tseng, Y.-H., Booth, I.R., and Saier, M.H. (2003). Two families of mechanosensitive channel proteins. Microbiology and molecular biology reviews: MMBR 67: 66-85-table of contents.

Rivera-Najera, L.Y., Saab-Rincón, G., Battaglia, M., Amero, C., Pulido, N.O., García-Hernández, E., Solórzano, R.M., Reyes, J.L., and Covarrubias, A.A. (2014). A Group 6 Late Embryogenesis Abundant Protein from Common Bean Is a Disordered Protein with Extended Helical Structure and Oligomer-forming Properties*. J Biol Chem 289: 31995–32009.

Urry, D.W., Hinners, T.A., and Masotti, L. (1970). Calculation of distorted circular dichroism curves for poly-l-glutamic acid suspensions. Arch Biochem Biophys 137: 214–221.

Urry, D.W. and Ji, T.H. (1968). Distortions in circular dichroism patterns of particulate (or membranous) systems. Arch Biochem Biophys 128: 802–807.

Uversky, V.N. (2019). Intrinsically Disordered Proteins and Their “Mysterious” (Meta)Physics. Aip Conf Proc 7: 10.

Uversky, V.N. (2013). The alphabet of intrinsic disorder: II. Various roles of glutamic acid in ordered and intrinsically disordered proteins. Intrinsically Disord Proteins 1: e24684.

Vacic, V., Uversky, V.N., Dunker, A.K., and Lonardi, S. (2007). Composition Profiler: a tool for discovery and visualization of amino acid composition differences. Bmc Bioinformatics 8: 211.

Veley, K.M., Maksaev, G., Frick, E.M., January, E., Kloepper, S.C., and Haswell, E.S. (2014). Arabidopsis MSL10 has a regulated cell death signaling activity that is separable from its mechanosensitive ion channel activity. The Plant cell 26: 3115–3131.

Verkest, C., Schaefer, I., Nees, T.A., Wang, N., Jegelka, J.M., Taberner, F.J., and Lechner, S.G. (2022). Intrinsically disordered intracellular domains control key features of the mechanically-gated ion channel PIEZO2. Nat Commun 13: 1365.

Wallmann, A. and Kesten, C. (2020). Common Functions of Disordered Proteins across Evolutionary Distant Organisms. Int J Mol Sci 21: 2105.

Wang, Y., Coomey, J., Miller, K., Jensen, G.S., and Haswell, E.S. (2021). Interactions between a mechanosensitive channel and cell wall integrity signaling influence pollen germination in Arabidopsis thaliana. J Exp Bot.

Wright, P.E. and Dyson, H.J. (2015). Intrinsically disordered proteins in cellular signalling and regulation. Nat Rev Mol Cell Bio 16: 18–29.

Yachdav, G. et al. (2014). PredictProtein—an open resource for online prediction of protein structural and functional features. Nucleic Acids Res 42: W337–W343.

Yang, J., Gao, M., Xiong, J., Su, Z., and Huang, Y. (2019). Features of molecular recognition of intrinsically disordered proteins via coupled folding and binding. Protein Sci 28: 1952–1965.

Yuan, F., Alimohamadi, H., Bakka, B., Trementozzi, A.N., Day, K.J., Fawzi, N.L., Rangamani, P., and Stachowiak, J.C. (2021). Membrane bending by protein phase separation. Proc National Acad Sci 118: e2017435118.

Zarin, T., Tsai, C.N., Ba, A.N.N., and Moses, A.M. (2017). Selection maintains signaling function of a highly diverged intrinsically disordered region. Proc National Acad Sci 114: E1450–E1459.

Zhou, H.-X., Rivas, G., and Minton, A.P. (2008). Macromolecular Crowding and Confinement: Biochemical, Biophysical, and Potential Physiological Consequences*. Annu Rev Biophys 37: 375–397.

