## Supplementary Figures for "The soluble N-termini of mechanosensitive ion channels MSL8, MSL9, and MSL10 are environmentally sensitive intrinsically disordered regions with distinct biophysical characteristics"

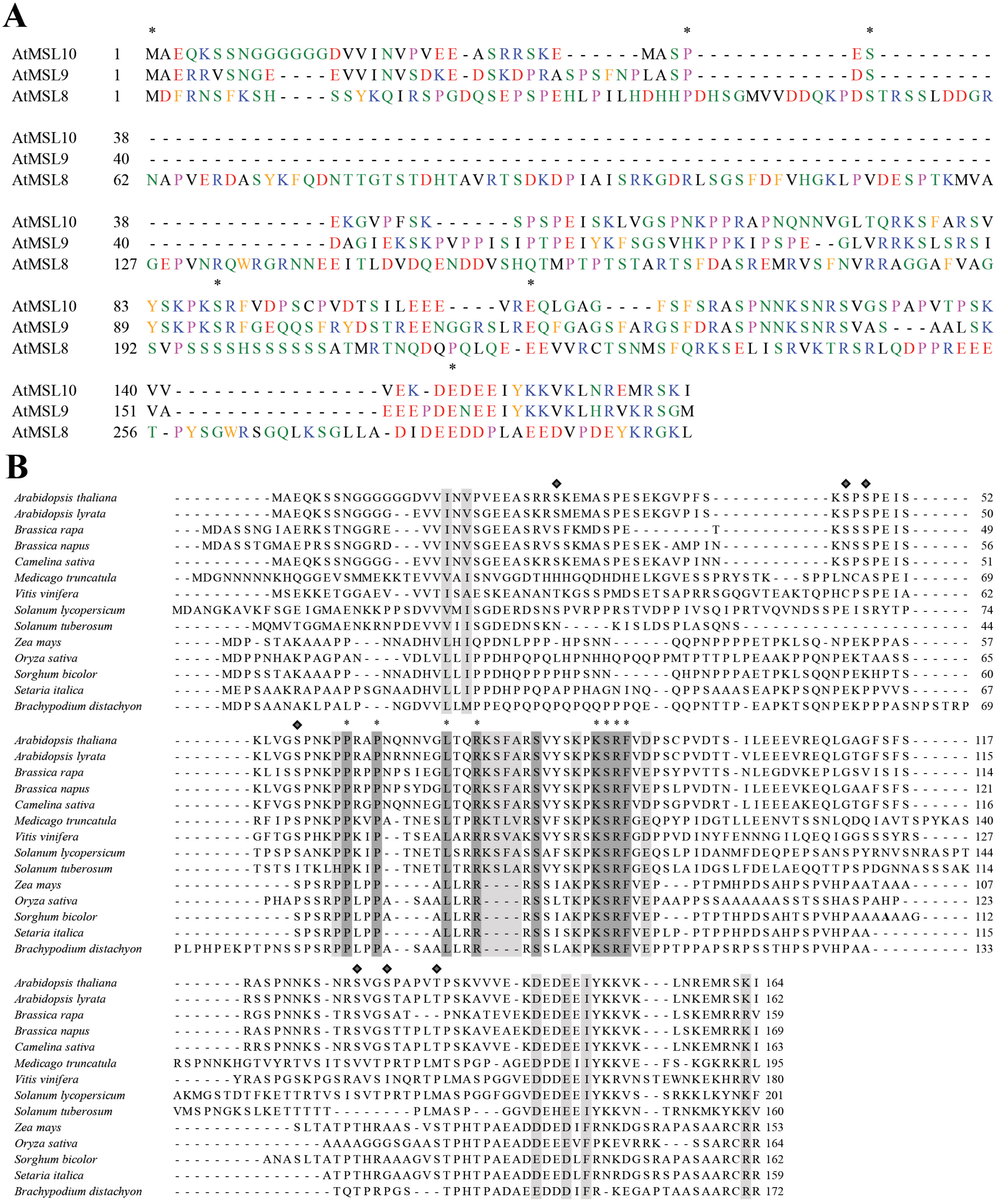


**Figure S1. The N-terminus of MSLs is hypervariable on the primary sequence level in plants.** (A) Alignment of *Arabidposis thaliana* MSL8, MSL9, and MSL10. Positions with identical residues are indicated with asterisks. Hydrophobic amino acids are in black, acidic residues in red, basic residues in blue, aromatic residues in orange, polar residues in green, and proline in pink. (B) Alignment of *Arabidopsis thaliana* MSL10 with 13 putative orthologs. Positions with identical residues are designated by darker shading and asterisks, while positions at which the properties of the amino acid are conserved are in lighter shading.

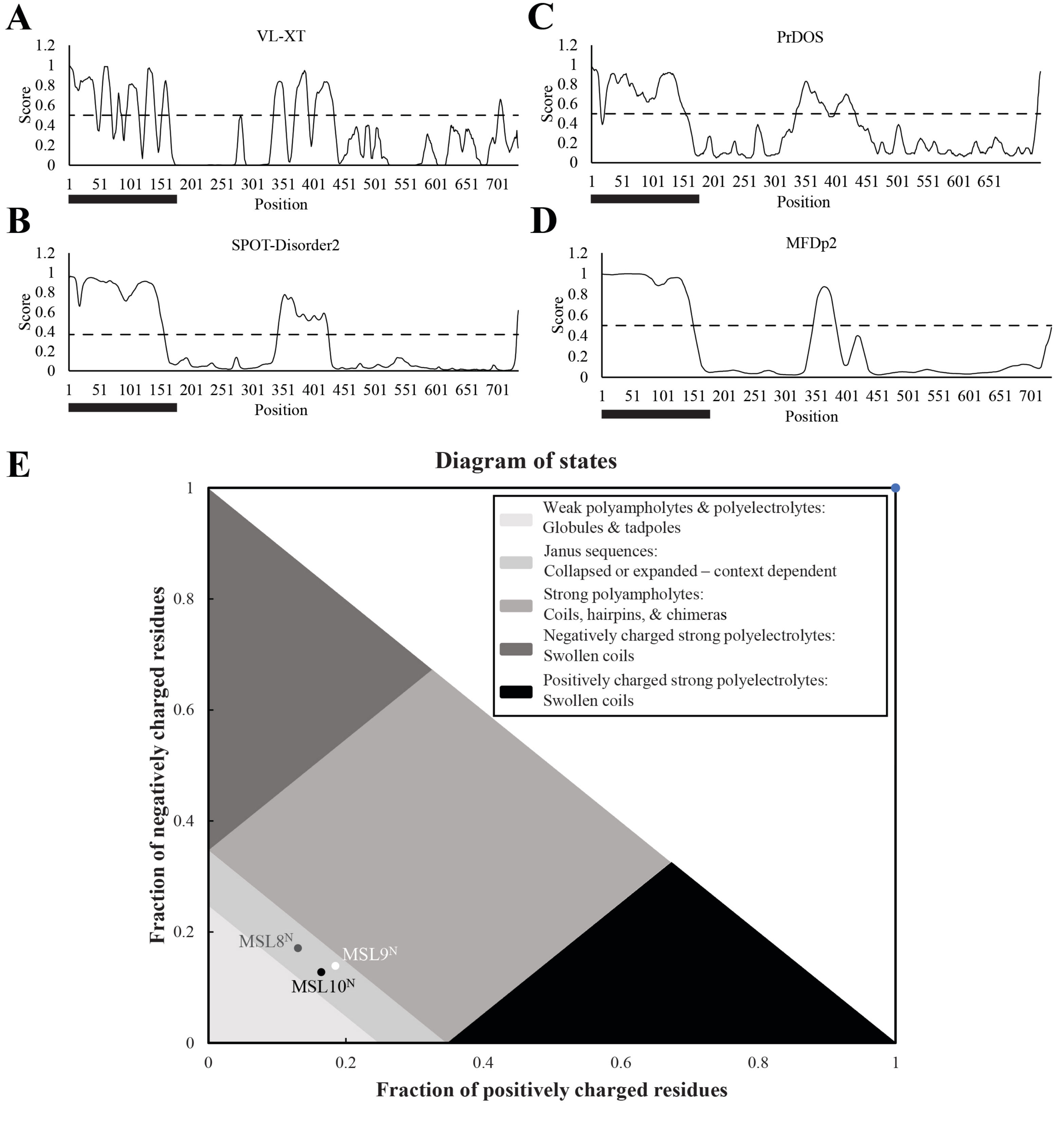


**Figure S2. The N-termini of MSLs are predicted to be disordered Janus sequences.** (A-D) Various disorder profiles for full-length Arabidopsis MSL10. Residues with a calculated disorder propensity greater than a given recommended algorithm threshold are predicted to be part of a disordered regions, indicated by the dashed black horizontal line. Black bar indicates the N-terminal residues of MSL10. (F) Positioning of $MSL8^{N}$, $MSL9^{N}$, and $MSL10^{N}$ on the Das-Pappu plot. The plot was generated using the CIDER webtool.

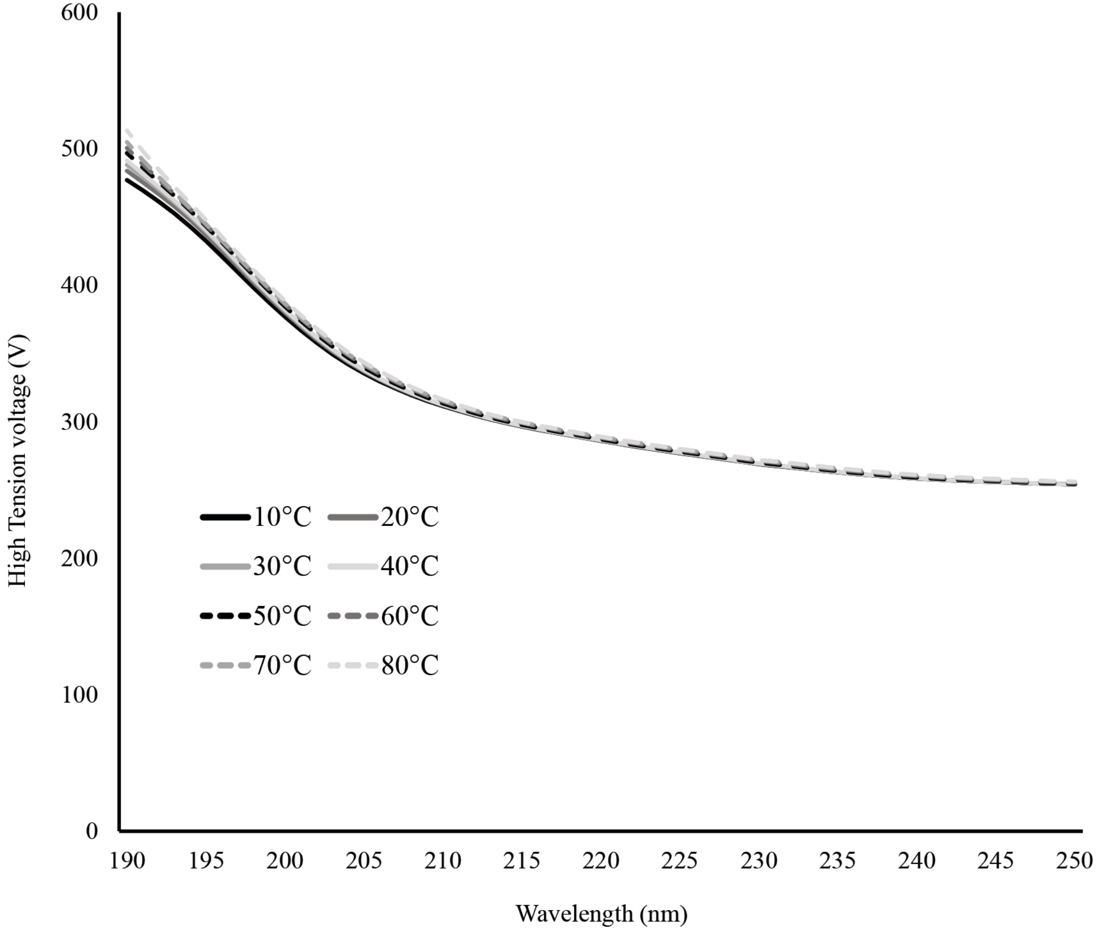


**Figure S3. High Tension voltage curves associated with MSL10$^{N}$ in increasing temperatures.** Measurements were obtained at 0.175 mg/mL protein in 20 mM sodium phosphate buffer, pH 7.4.

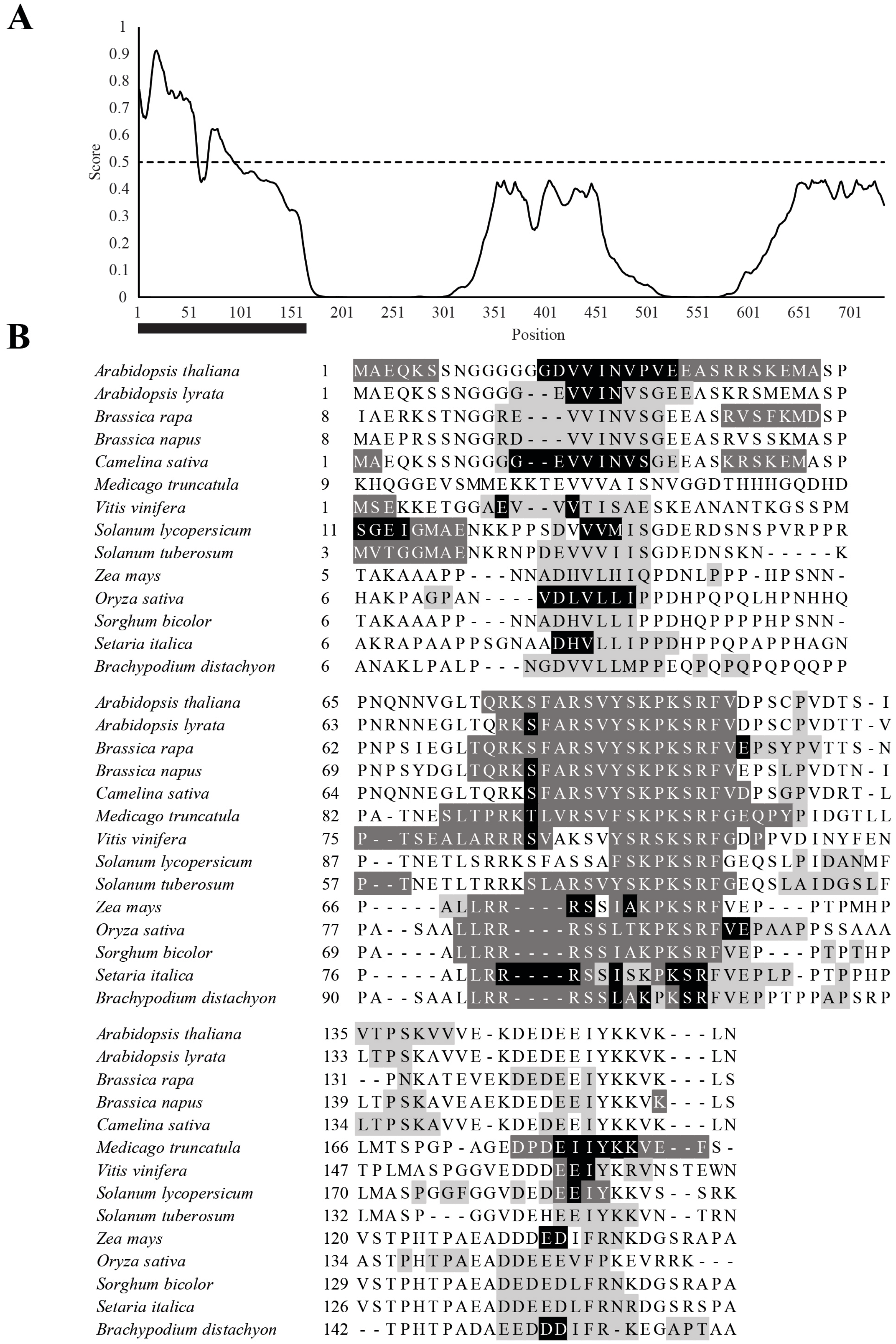


**Figure S4. Predicted sites of protein interaction in the N-terminus of MSL10 and putative orthologs.** (A) ANCHOR2 profile for full-length MSL10. Sequences assigned a score greater than 0.5 are predicted to be protein binding regions, as indicated by the dashed black horizontal line. The black bar indicates the residues of the MSL10 N-terminus. (B) MoRF regions identified by the MoRFPred and MoRFChibi_Web webservers for *Arabidopsis thaliana* MSL10 and 13 putative MSL10 orthologs. Amino acids predicted by MoRFPred (P > 0.5) are highlighted in light grey and MoRFChibi_Web (MCW > 0.7) predictions are highlighted in dark grey. Amino acids identified as potential MoRF residues by both MoRFPred and MoRFChibi_Web are highlighted in black.
